# How does target lesion selection affect RECIST? A computer simulation study

**DOI:** 10.1101/2022.04.14.488203

**Authors:** Teresa T. Bucho, Renaud Tissier, Kevin Groot Lipman, Zuhir Bodalal, Andrea Delli Pizzi, Thi Dan Linh Nguyen-Kim, Regina Beets-Tan, Stefano Trebeschi

## Abstract

RECIST is grounded on the assumption that target lesion selection is objective and representative of the change in total tumor burden during therapy. A computer simulation model was designed to challenge this assumption, focusing on a particular aspect of subjectivity: target lesion selection. Disagreement among readers, and between readers and total tumor burden was analyzed, as a function of the total number of lesions, affected organs, and lesion growth. Disagreement aggravates when the number of lesions increases, when lesions are concentrated on few organs, and when lesion growth borders the thresholds of progressive disease and partial response. An intrinsic methodological error is observed in the estimation of total tumor burden (TTB) via RECIST. In a metastatic setting, RECIST displays a non-linear, unpredictable behavior. Our results demonstrate that RECIST can deliver an accurate estimate of total tumor burden in localized disease, but fails in cases of distal metastases and multiple organ involvement. This is worsened by the “selection of the largest lesions”, which introduce a bias that makes it hardly possible to perform an accurate estimate of the total tumor burden. Including more (if not all) lesions in the quantitative analysis of tumor burden is desirable.

## 1 INTRODUCTION

The Response Evaluation Criteria in Solid Tumors (RECIST) consists of a standardized methodology used in clinical trials to evaluate tumors’ response to therapy (Seymour et al. 2010), by defining endpoints surrogate for overall survival, namely progression-free survival and overall response rate (Villaruz and Socinski 2013; Mushti, Mulkey, and Sridhara 2018). The most recent version of the criteria, RECIST 1.1 (Eisenhauer et al. 2009), states that a maximum of five measurable lesions, and no more than two per organ, should be selected as target lesions and measured at baseline. On follow-up, the same target lesions should be identified and re-measured. Response to therapy is then classified into four categories, primarily based on the percent change of the sum of the sizes of target lesions between baseline (or at nadir) and follow-ups.

Target lesion selection assumes that target lesions can be objectively and reproducibly identified and measured (Villaruz and Socinski 2013). However, inter- and intra-variability when measuring tumor size, and identification of new lesions are known factors contributing to inconsistency in the application of RECIST (Skougaard et al. 2012; Abramson et al. 2015; Sridhara, Mandrekar, and Dodd 2013; Beaumont et al. 2018).

Particularly, disagreement in the selection of target lesions is one of the leading causes of variability in RECIST (Tovoli et al. 2018; Keil et al. 2014; Kuhl et al. 2019; Iannessi et al. 2021). According to the guidelines, the largest lesions in diameter, or the ones that “lend themselves to reproducible repeated measurements” should be chosen (Eisenhauer et al. 2009). However, the interpretation of these guidelines is reader-dependent (Keil et al. 2014), and different readers might end up selecting different target lesions, even if correctly applying RECIST criteria. In other words, even when excluding the factor of diameter-measuring error (e.g. through medical segmentation software), RECIST still shows intrinsic variability if a limited number of target lesions has to be selected.

RECIST strives to make the comparison of clinical trial outcomes possible and reproducible (Villaruz and Socinski 2013). However, this is grounded on the assumption that the target lesions are objectively identified and that this subset of the total tumor burden (TTB) of a patient is sufficient to adequately represent response to therapy (Schwartz et al. 2003; Keil et al. 2014; Kuhl et al. 2019). Nevertheless, these assumptions may not hold in practice.

In this study, we aim to show that target lesion selection introduces large inconsistency in RECIST assessments and that its role as a surrogate of TTB is not sustained. Motivated by a previous study by Moskowitz et al. (Moskowitz et al. 2009), we created a computer simulation model to investigate target lesion selection variability in RECIST across different patient characteristics, and its validity as a proxy for TTB. Compared to observational studies, where we are limited to the characteristics of the retrospective cohort (e.g. number of patients, lesions per patient, average tumor growth, unavailability of total tumor burden measurements), a simulation model yields advantages. It allows us to run a controlled experiment, in which we can generate any virtual cohort of patients with precise characteristics, and with which we can study the influence of each of them on the outcome: the RECIST assessment.

## 2 METHODS

### 2.1 Simulation Model

We created a computer simulation model, represented schematically in Figure 1. With this model, we generate cohorts of patients undergoing a virtual clinical trial. A cohort of patients is characterized by four parameters: maximum number of lesions (L_max_), maximum number of organs involved (O_max_), mean tumor percent growth (μ), and growth variance (Σ). To test the influence of each of the parameters on the RECIST assessment, we fix, in succession, all but one of the four input parameters to a default value, while the remaining variable changes within a certain range: L_max_ between 1 and 20, O_max_ between 1 and 10, μ between −100% and 200%; and μ between 10 and 100%. Σ is composed of three different variances, of which two change (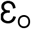 and 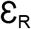, see next section), with values ranging between 10^2^ and 100^2^ ‰, and one is kept at its default value (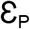, see section 2.2 and Supplement 1).

**Figure 1:**
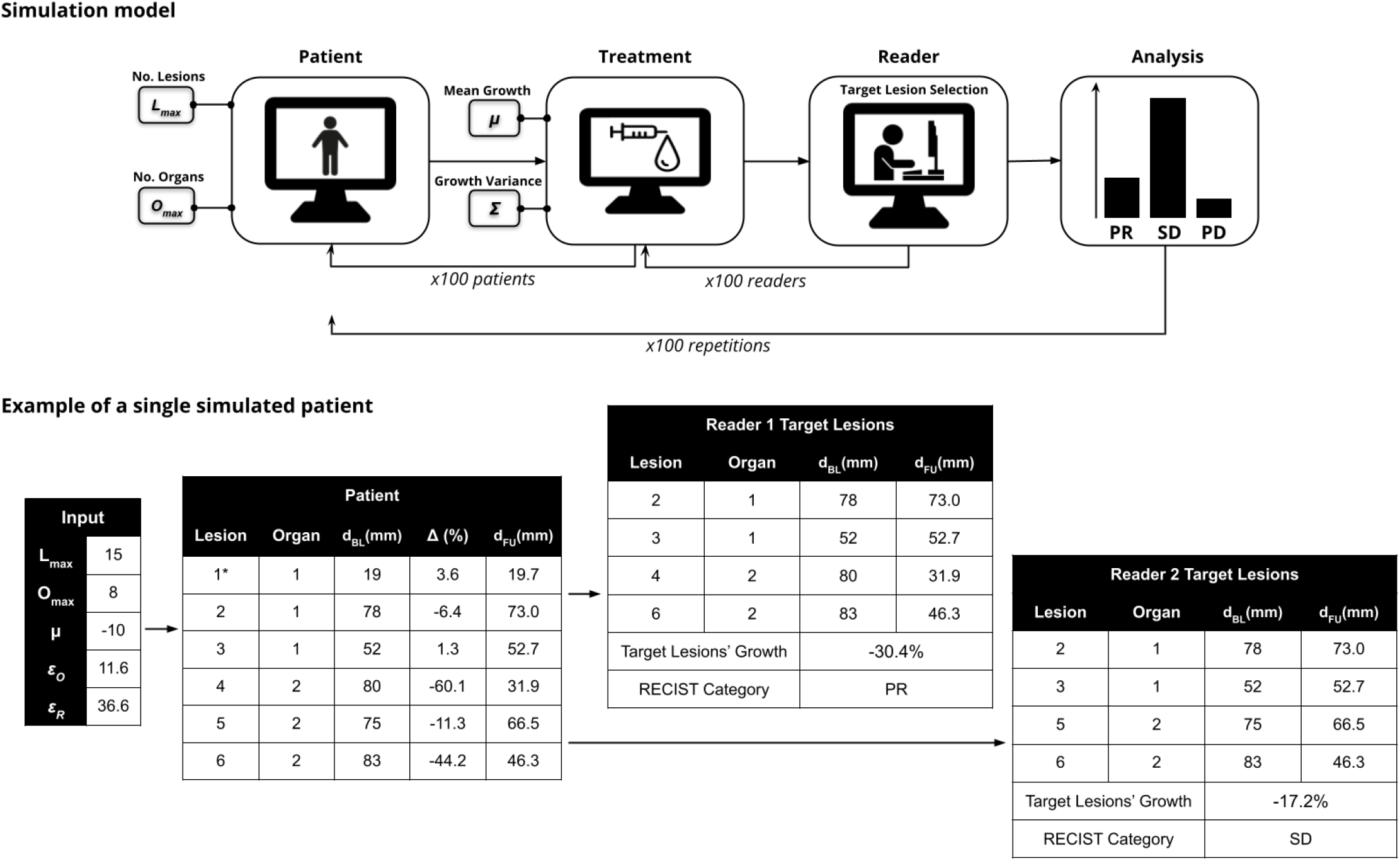
Top: schematic representation of the simulation model. Number of lesions (L) and number of organs (O) are, for each patient, a random number between [1, L_max_] and [1, O_max_], respectively. Each lesion’s percent growth (Δ) is sampled from a truncated multivariate normal distribution with mean μ and covariance matrix Σ (a combination of 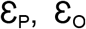, and 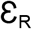). Bottom: example of a simulated patient with L=6 lesions distributed among O=2 organs, and two virtual readers. Only large lesions are eligible for selection as target lesions, so lesion 1(*) is excluded. The two readers disagree on the choice of target lesions, resulting in different RECIST categories.

### 2.2 Virtual Patient

Each simulated patient can be described in four steps: 1) the number of organs (O) and lesions (L) of each patient is drawn randomly between [1 to O_max_] and [1 to L_max_] respectively. The lesions are distributed among the organs with a multinomial distribution, where the probability of a lesion getting assigned in each organ is the same (if O > L then some organs get assigned no lesions); 2) for each lesion, a random baseline tumor size (d_BL_) between 10 and 100 mm is sampled from a random normal distribution; 3) the growth or shrinkage of each lesion (Δ) is simulated via a truncated multivariate normal distribution with mean μ and variance-covariance matrix Σ. Σ is composed of 3 different variance parameters to account for the growth correlation between lesions within patients 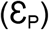, within organs 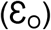, and the residual variance 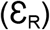; 4) based on the sampled percent growth, Δ, and the baseline diameter d_BL_ of each lesion, we calculate the follow-up diameter (d_FU_). We only simulate these two time points of the virtual clinical trial: this will suffice to explore the effects of RECIST variability on the estimation of the TTB without overcomplicating the simulation unnecessarily. An example of a simulated patient can be seen in Figure 1.

The default values of 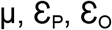, and 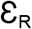 were estimated from three real datasets: n=61 patients with melanoma treated with immunotherapy, n=44 urothelial cancer patients treated with immunotherapy, and n=37 patients with non-small cell lung cancer treated with chemotherapy, already reported in previous work (S Trebeschi et al. 2019; Stefano Trebeschi et al. 2021). 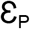 has no impact on target lesion selection, and thus its impact on variability was not analyzed. For all datasets, the diameters of all lesions present at both baseline and follow-up were available. From that, the respective growth percent was estimated, on which linear mixed-effects model was fit. Lmax and O_max_ were set to 3 different pairs of values (5, 2), (10, 4), and (15, 8), aiming to portray different stages of disease — low, medium, and high, respectively.

### 2.3 Virtual Reader

Each patient was evaluated by 100 independent virtual readers. A virtual reader represents a radiologist that selects up to 5 target lesions, no more than 2 per organ, according to the RECIST criteria. To account for the RECIST guidelines suggesting that the largest lesions should be selected as target lesions, we restricted the choice of target lesions to a pool of lesions with size within 20% of the size of the largest lesion in each organ. When no lesions were under this condition, we added the second largest lesion to the pool, as well as all the lesions within 20% of its size (thus allowing at least 2 lesions to be chosen per organ). For each patient, we calculate the overall percent growth based on the sum of diameters of target lesions selected by each reader and assign the respective RECIST category. A percent growth above 20% (and minimum absolute increase of 5 mm) corresponds to progressive disease (PD), below −30% to partial response (PR), equal to −100% to complete response (CR), and otherwise, stable disease (SD) (Eisenhauer et al. 2009).

### 2.4 Data analysis

To study the relationship between readers’ disagreement and patients’ characteristics, we simply record the percentage of patients with inconsistent RECIST readings across different readers for a specific set of cohort characteristics (i.e. model parameters). To minimize the effect of random noise on the analysis, each experiment is repeated and averaged over 100 runs. The disagreement levels for different combinations of patient cohort characteristics and treatment effects are recorded and analyzed.

To study the accuracy of RECIST as an estimator of the changes in the TTB, we turn the analysis into a classification problem. Given that, for each simulated patient, we know the true change in TTB (and therefore the true response class of PD, SD, or PR) we simply count the number of RECIST readings that resulted in an erroneous prediction of TTB-derived class of response. As in the analysis above, we repeat and average over 100 runs to minimize the effect of random noise, and study the trend of RECIST misclassification of TTB as a function of the number of lesions (L_max_ = L). Figure 2 shows a schematic representation of the analysis. The code of the simulation model is publicly available on our GitHub repository^1^.

**Figure 2:**
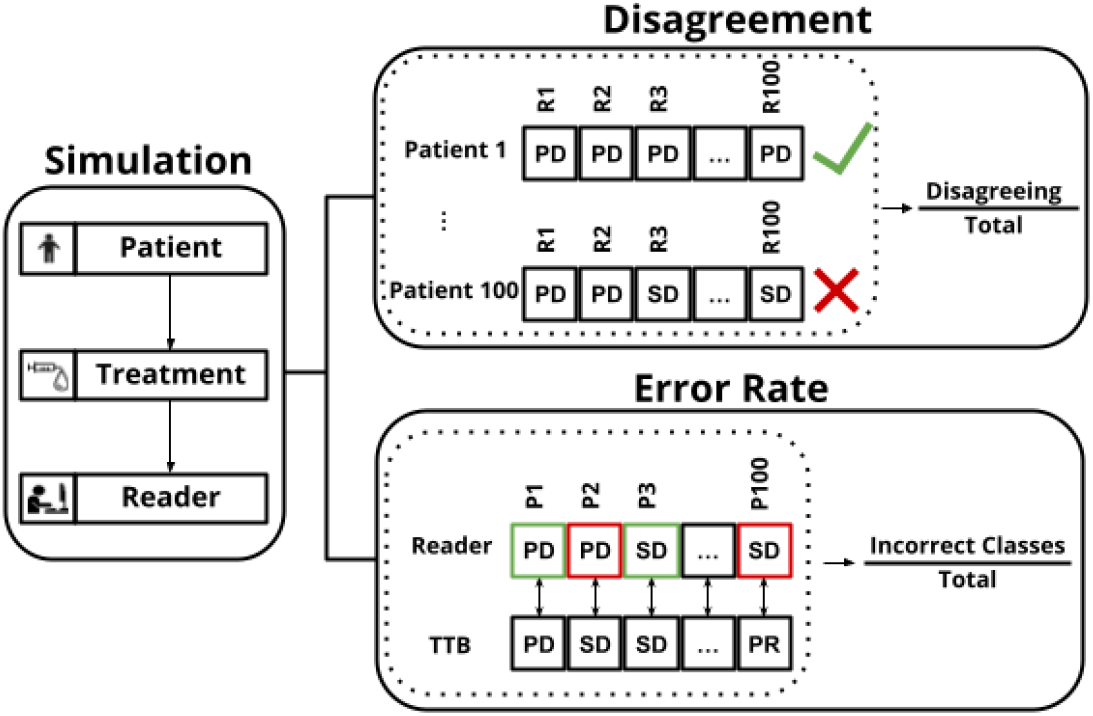
Metrics for analyzing RECIST variability in the simulated cohorts of patients. Disagreement corresponds to the percentage of patients in a cohort with inconsistent RECIST readings (i.e. different response categories attributed by different readers). Disagreement is then analyzed as a function of the input parameters (L_max_, O_max_, μ). The error rate, computed per reader, corresponds to the percentage of incorrectly attributed response categories, taking TTB as reference. The error rate is analyzed as a function of the L_max_, and for different variations of RECIST (see results section). R = Reader; P = Patient.

## 3 RESULTS

We first investigate disagreement between readers as a function of patient characteristics. Figure 3 depicts RECIST variability across different cohort characteristics. We can observe a linear relation between the maximum number of lesions per patient (L_max_) and disagreement, which behaves independently from the extent of the disease spread (Figure 3). Even when the maximum number of lesions is less than five, we still observe some disagreement, since only a maximum of two lesions per organ can be chosen.

**Figure 3:**
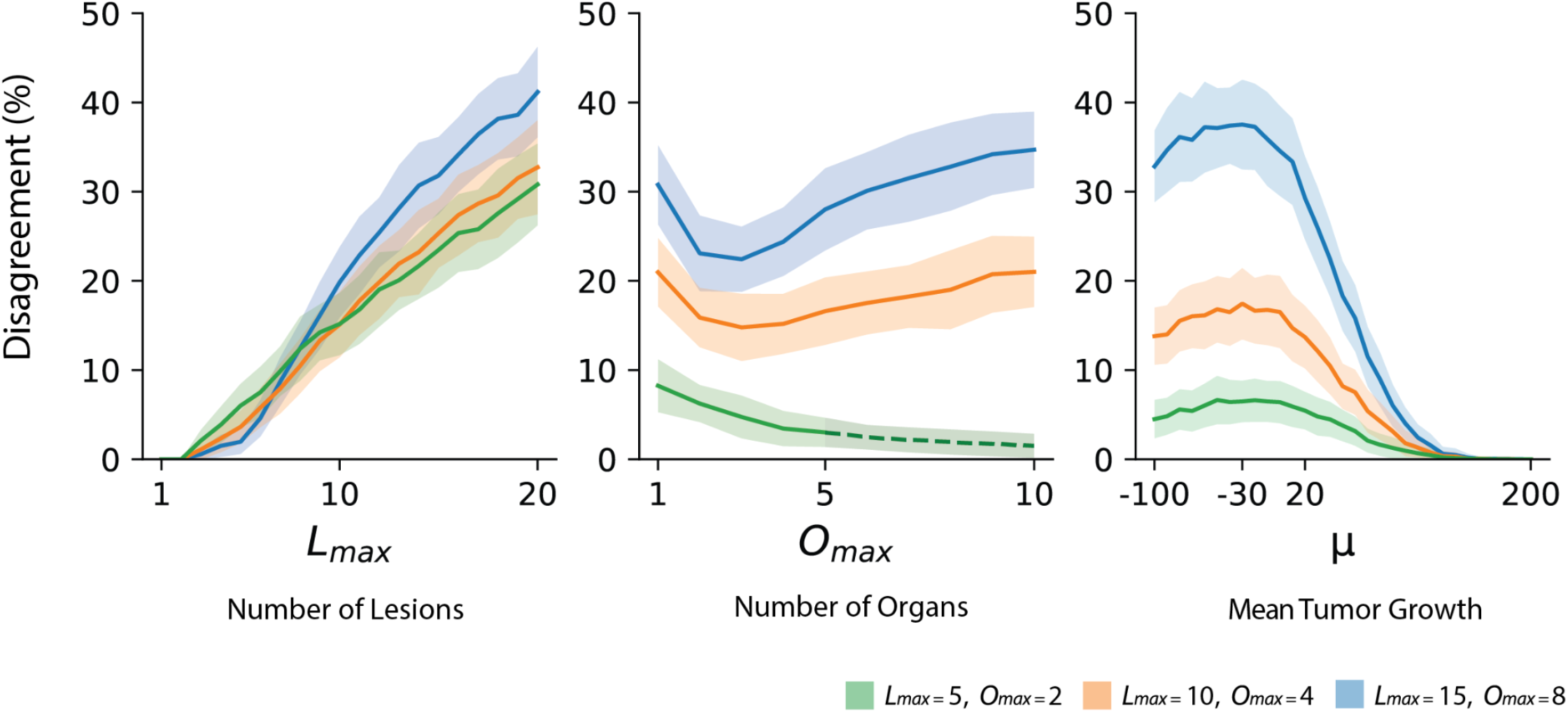
Mean disagreement levels as a function of the number of lesions, number of affected organs, mean lesion growth, with respective confidence intervals (± standard deviation). The different curves represent different combinations of the default values of L_max_ and O_max_, respectively: blue: (5, 2); red: (10, 4); and black (15, 8), representing different stages of disease. In plots A and B, only the default values of O_max_ and L_max_, respectively, change. Dashed line: when O_max_ > L_max_, disagreement decreases as the likelihood of the lesions becoming more spread out increases.

There is a nonlinear relation between disagreement levels and the number of affected organs (O_max_). When O_max_ varies from one to ten, we observe high disagreement initially, since many lesions are concentrated on very few organs and due to the imposition of a maximum of two lesions per organ, readers are forced to make different choices of target lesions. Disagreement then decreases since lesions spread out enough over multiple organs, limiting the number of choices, but increases again due to the limit of a maximum of two target lesions per organ, forcing a choice of target lesions and, implicitly, target organs. When Lmax = 5 (green line in the middle plot of Figure 3), disagreement approaches zero when Omax increases, since all lesions can be selected if no organ contains more than two lesions (Figure 3). When Omax increases beyond the number of lesions, some organs do not get assigned any lesions. We still observe that disagreement continues to decrease merely because the chances of the 5 lesions being located in different organs increases (indicated by the dashed line). This scenario, where O_max_ > L_max_, can also occur in other curves but to a negligible degree.

Disagreement among readers is higher when the average tumor growth (μ) borders the thresholds of PD and PR (+20% and −30% respectively). When distancing from the thresholds, disagreement decreases since it becomes more likely for either PD or PR to be attributed. A nonlinear relation exists between average tumor growth and disagreement which, looking across the different groups of disease stages, seems to be amplified by the stage or spread of the disease. The disagreement levels as a function of the input variances can be found in Supplement 2.

We then investigated the performance of RECIST to predict the true class of response, as defined by the TTB, as a function of the number of lesions. In figure 4A, it can be observed that the error rate plateaus above 15% for all patients with 10+ lesions, and reaches levels around 20% for patients with 20 lesions, for all levels of Omax. For comparison, we re-ran the experiments with different variations of RECIST: RECIST 1.0, which allows the selection of up to ten target lesions; and RECIST-random, where the readers are free to choose any target lesions, and are not limited by the largest ones (Figure 4B). RECIST 1.0 reaches the lowest error rate, staying around 10%, even for patients with 20 lesions. Interestingly, both RECIST-random 1.1 and RECIST-random 1.0 perform only slightly worse than the original counterparts, with RECIST-random 1.0 achieving a better estimation of the true response class than standard RECIST 1.1.

**Figure 4:**
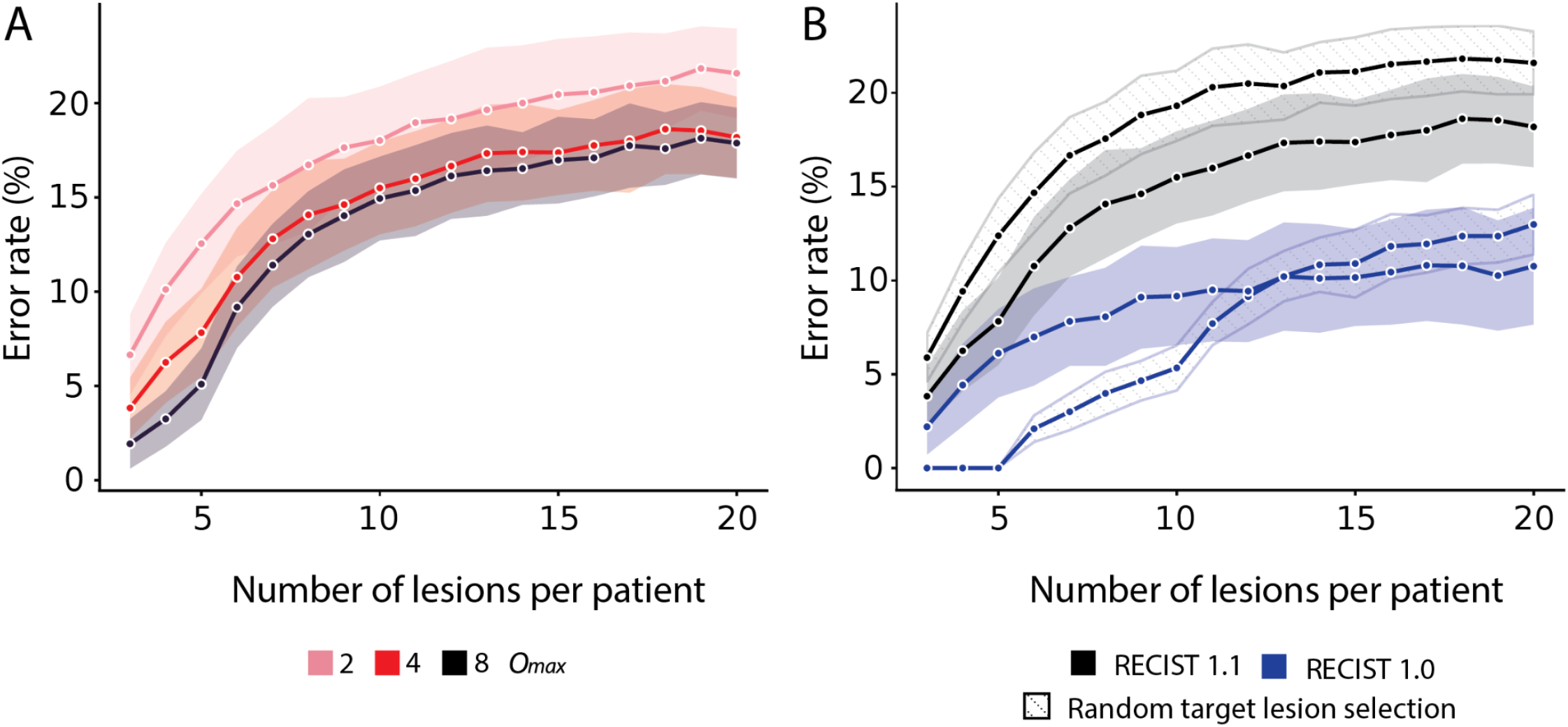
A) Error rate in classifying response with RECIST compared to the true tumor response, as defined by TTB, for different values of O_max_ B) Error rate for O_max_ = 4, comparing variations of RECIST.

## 4 DISCUSSION

This study aimed to investigate target lesion selection as a cause of variability in RECIST, in a controlled experiment, by means of simulated models. Previous studies have identified and studied target lesion selection as a source of variability in response classification in real cohorts of patients (Keil et al. 2014; Tovoli et al. 2018; Kuhl et al. 2019). The advantage of a controlled experiment over an observational study is that it allows us to set the effects of the treatment and the characteristics of the patient cohort, and analyze the particular conditions under which RECIST is inconsistent, despite being correctly employed. This helps us understand where the focus of future developments in tumor size-based response assessment should be: in polishing the already existing RECIST guidelines or in searching for better alternatives. Overall, our findings suggest a complex function linking patient characteristics with RECIST variability. In other words, while it is possible to explain the behavior of RECIST variability in relation to different characteristics of the patient cohorts, it seems impossible to draw a general rule that defines the expected level or behavior of RECIST variability in a simple fashion.

We observed that disagreement between readers increases linearly (and nearly independently from the number of organs) when the total number of lesions per patient increases, since the chances of different readers selecting the exact same target lesions decreases. RECIST recommends choosing target lesions based on size or on how reproducible their measurement is, which should help dilute differences between readers and lead to reduced variability. Selecting the largest lesions in diameter implicitly requires that lesions are sufficiently distinct in size such that the ones with the largest diameters are easily and uniquely identified, or that actual measurements of all the lesions are available such that the largest can be objectively chosen (Moskowitz et al. 2009). Even when same-lesion actual measurements are carried out, there can be critical inter- and intra-observer measurement discrepancies (Moertel and Hanley 1976; Skougaard et al. 2012; Yoon et al. 2016; Tovoli et al. 2018). Therefore, if the selection of the largest lesions has to be performed by visual inspection it is expected that disagreements will aggravate further (Moskowitz et al. 2009), especially if the lesions have complex shapes. In our simulation, we allowed the virtual readers to select lesions from a pool of “large lesions”, which we set to be all those within 20% of the size of the largest lesion, to account for the possibility of readers diverging in their discernment of what the largest lesions should be. Selecting the most reproducible lesions is also heavily subject to personal judgment. Readers need to decide if a lesion is “reproducible enough” or even of malignant nature (Schwartz et al. 2003; Iannessi et al. 2021) and if it should be chosen in favor of a larger lesion. Furthermore, selecting the largest or the lesions with the most well-defined boundaries for measurement may imply that the lesions selected are the ones responding to treatment in a similar way (Kuhl 2019), while largely overlooking the (non-target) lesions that could be as or more determinant for treatment response assessment (Coy et al. 2019). Despite the criteria guiding the choice of target lesions, this is one of the main contributors to RECIST’s inconsistency (Tovoli et al. 2018; Keil et al. 2014; Kuhl et al. 2019). As noted by (Kuhl 2019), while additional recommendations would indeed help reproducibility, these might also just be masking the fundamental problems of RECIST and constraining readers from selecting the same lesions, even if these are not representative of the true tumor burden.

The substantial disagreement in response assessment by different readers suggests that 5 lesions alone are not sufficient to assure an objective response assessment. This is confirmed in our experiments by the relatively high error rate of RECIST in predicting the true response class (based on the TTB), in comparison to the substantially lower error rate reported for RECIST 1.0 (which allows double the number of target lesions in the analysis) and the relatively small difference in performance with an alternative, hypothetical RECIST which allows the selection of random target lesions, instead of the largest ones. Seeing that the selection of random lesions (instead of the largest ones) did not degrade the performance as much as decreasing the number of target lesions (from ten to five) suggests that the inclusion of the largest lesions had a small effect. Including more, if not all lesions in the quantitative analysis would be the best way to reduce variability and help capture the true TTB, likely leading to a more representative evaluation of response to therapy (Keil et al. 2014; Kuhl 2019), and possibly fewer patients required in clinical trials to prove the efficacy of the treatment. Since percent change is computed by taking into account the sum of the sizes of all target lesions, more importance is given to changes in larger lesions. This explains why we observe an increase in error when selecting random lesions compared to only large ones, which is aggravated if the number of target lesions is more limited.

In the simulation study of Moskowitz et al. 2009 (Moskowitz et al. 2009), the impact of the number of target lesions measured (ten, five, three, two, or one) on response assessment was investigated. The authors agreed that measuring a smaller number of lesions led to a larger percentage of misclassified patients. Nevertheless, it was concluded that measuring five lesions was the best compromise between a good enough proxy for TTB and the labor-intensive task of assessing many lesions, since assessing ten lesions did not provide added benefit. In (Schwartz et al. 2003), the authors defend that the number of lesions to be measured should be established based on the specific context of the study and intended comparisons with other studies.

In some therapies, clinical outcomes are associated with the site of metastasis (Bianchi et al. 2020; Lu et al. 2019; Schmid et al. 2018). As patients usually have more than one site of measurable disease, spreading the choice of lesions across the body (by imposing a maximum of 2 target lesions per organ) allows for the selection of a complete set of lesions with varying degrees of response to therapy. Nevertheless, as we observed, when many lesions are concentrated in a single or few organs, this imposition creates high disagreement between readers. Including all measurable lesions together with a subanalysis of the tumor burden per organ could be warranted.

Disagreement between readers varied non-linearly with the average tumor growth but increased linearly with disease stage. It was aggravated when the mean growth of single lesions was centered around the small percentages of growth or shrinkage. The further away mean lesion growth of the set of target lesions is from the cutoffs for progressive disease (20%) and partial response (−30%), the clearer the attribution of the response category is. However, when mean lesion growth is closer to these cutoffs, classifying response is more troublesome and completely dependent on the selected lesions. The coarse compartmentalization of response into four categories has been the target of critique. It has been proposed that a system that describes lesion growth as a continuous variable (Moskowitz et al. 2009; Kuhl 2019; Mercier et al. 2019), would be more appropriate for assessing response to therapy (Jain et al. 2012; Wang et al. 2019). Furthermore, while percent change allows us to compare patients with different levels of baseline tumor burden easily, one could also question the appropriateness of this metric. For example, should the doubling in size of a lesion with a very small diameter at baseline be considered as relevant for response assessment as the doubling in size of a lesion with a much larger baseline size? Time, estimated as a rate of growth, should also be accounted for.

Nevertheless, our results must be interpreted in the light of the default values chosen for the input parameters and the overall behavior of the simulation curves should be the focus point. We estimated the default values of mean lesion growth and variance from three real datasets of patients, as a way of approximating two possible scenarios of growth patterns between baseline and first follow-up. For example, if patient, organ, and residual default variances were very small, we would expect to see two disagreement peaks centered around the exact cutoff percentages for PD and PR. Due to the nature of the simulation, where lesion growth is sampled from a truncated multivariate normal distribution with non-small variance, we do not observe CR, and we still observe some disagreement when μ = −100%. In our simulation model, we did not explicitly take into account selection variability arising from the identification of what should be considered “reproducible” lesions. The readers were simulated such that they had to choose a maximum of five and no more than two lesions per organ, and only from the largest pool of lesions. However, there was no other explicit rule to force the readers to spread their choice of lesions as much as possible through the organs. Reader experience, non-target lesion qualitative assessment, and possible appearance of new lesions were also not considered.

Volumetric measures have been described as a better discriminator of tumor size changes than RECIST’s linear measurements, related to the capacity of tumor volumetry better describing the size of irregular lesions (Fenerty et al. 2016; Oubel et al. 2015) and to the reduced interobserver variability (Rothe et al. 2013; Wulff et al. 2013; Zimmermann et al. 2021). While manual tumor segmentation is still very labor-demanding, advances in artificial intelligence for automatic volumetric tumor segmentation of whole-body scans (Jemaa et al. 2020; Tang et al. 2021) might soon be a solution in clinical practice. Such a development could change the landscape of response assessment by replacing target lesion selection with TTB estimation and rethinking the coarse threshold-based compartmentalization of response.

In conclusion, RECIST’s response assessment suffers from large variability due to the selection of target lesions, especially in a metastatic setting. Even if readers were to agree on the same set of target lesions, it cannot be guaranteed that these are representative of TTB and response to treatment. Including more, if not all, measurable lesions in the quantitative assessment is desirable, possibly aided by automatic segmentation models.

## SUPPLEMENT 1 - Default values for input parameters

In this table, the default values for 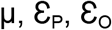 and 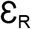, as estimated by the linear mixed-effects model fitted on the three real datasets (N=61 patients with melanoma, N=44 urothelial cancer patients and N=37 patients with non-small cell lung cancer) are listed.

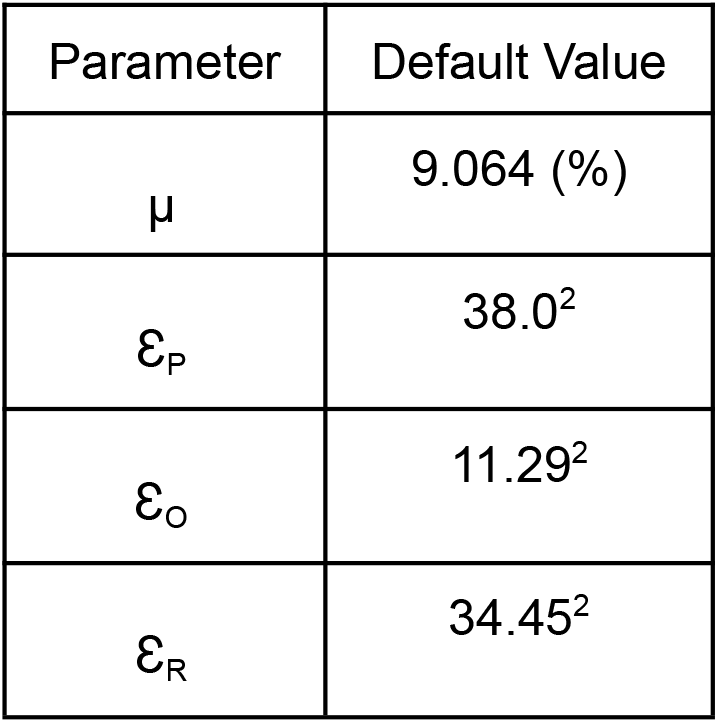

| Parameter | Default Value |
| --- | --- |
| $\mu$ | 9.064 (%) |
| $\mathcal{E}_P$ | 38.0<sup>2</sup> |
| $\mathcal{E}_O$ | 11.29<sup>2</sup> |
| $\mathcal{E}_R$ | 34.45<sup>2</sup> |

## SUPPLEMENT 2 - Variance simulation plots

The disagreement between readers seems robust to variance between-organs 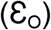 and only decreases for large values, which could be due to the use of a truncated multivariate distribution. When the residual variance 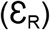 increases we observe that the disagreement between readers increases. This is expected as the residual variance represents variations between lesions independent of the fact that these lesions are from the same patient or the same organs. This means that with a large residual variance, lesions within the same organ of the same patient will tend to respond completely differently to the treatment.

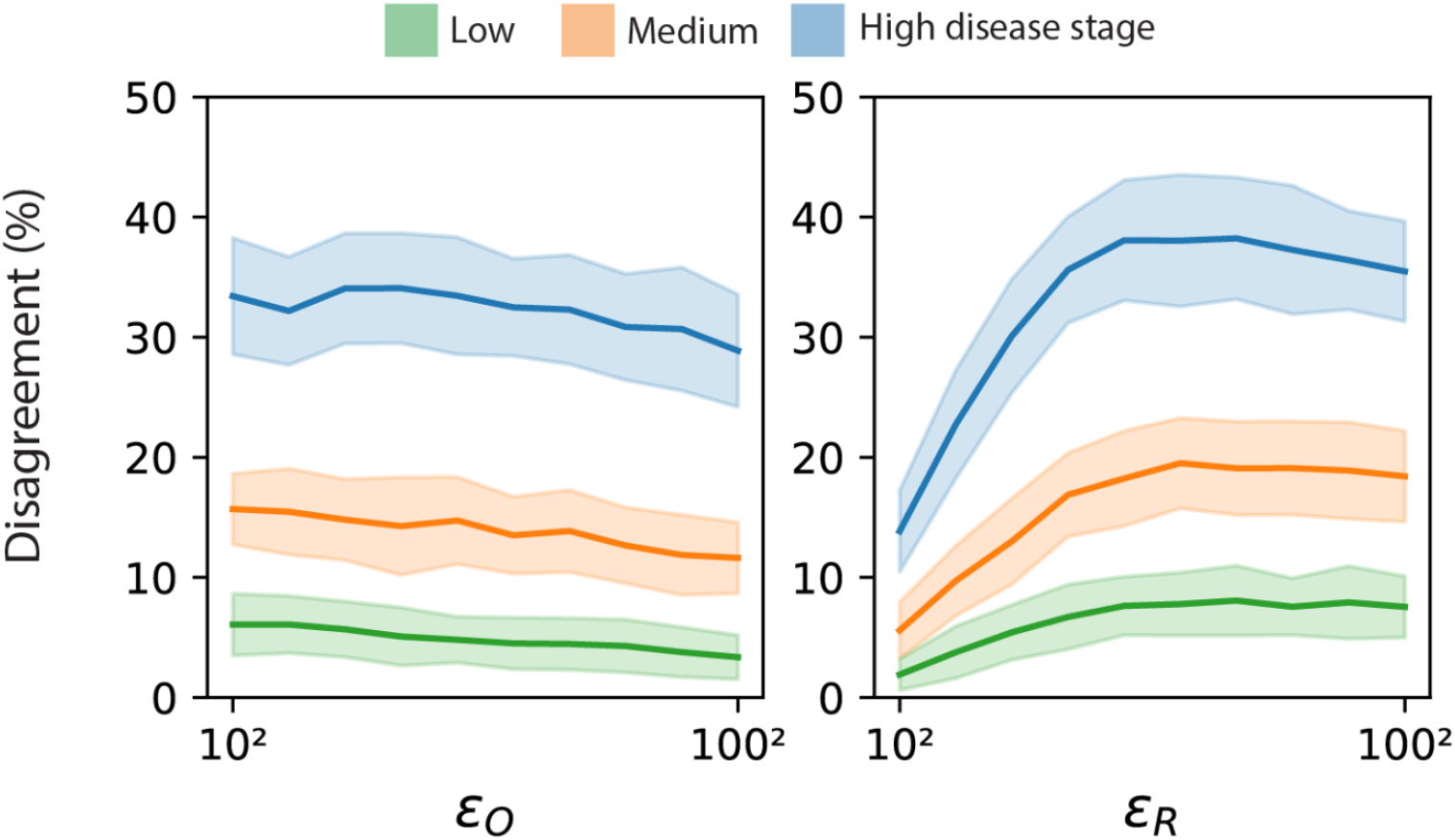

## SUPPLEMENT 3 - Error rates for O_max_ = 2 and O_max_ = 8

Performance of RECIST to predict the true class of response, as defined by the TTB, as a function of the number of lesions, for O_max_ = 2 and O_max_ = 8.

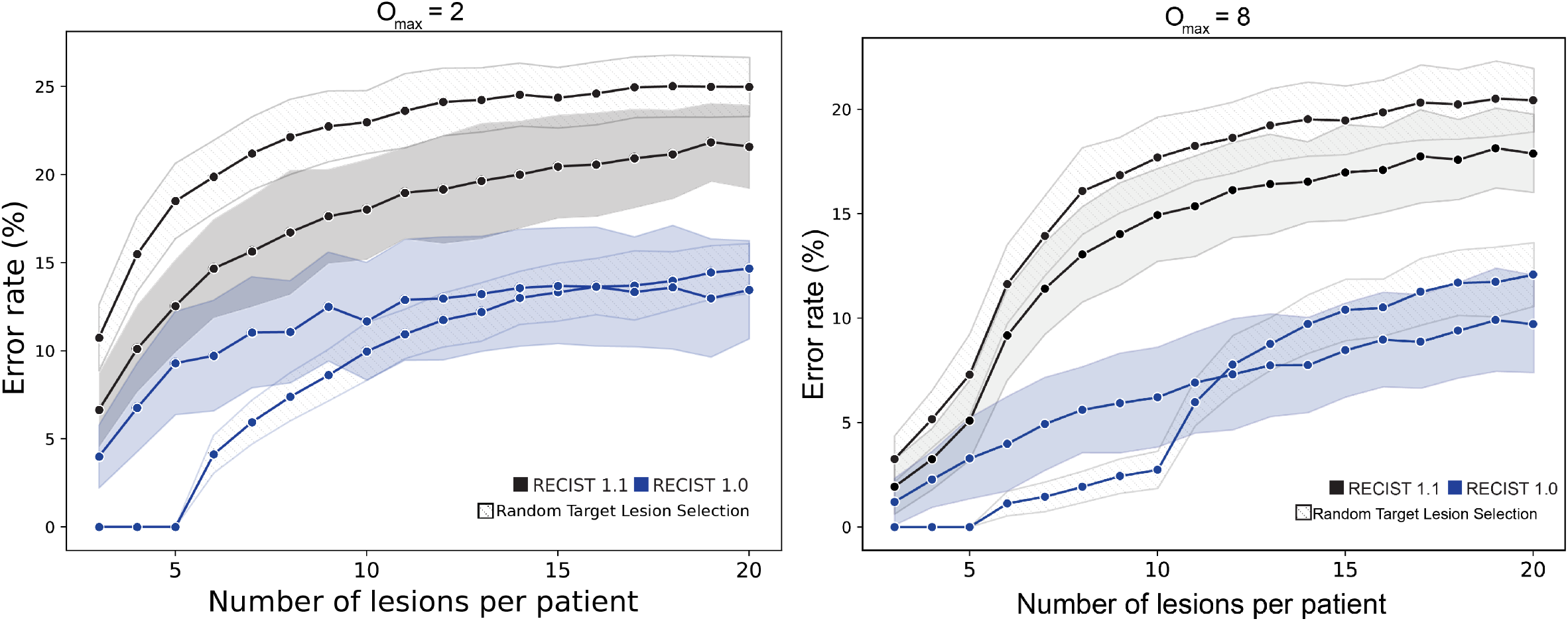

## Footnotes

1 https://github.com/nki-radiology/recist-variability

